# The predictive power of intrinsic timescale during the perceptual decision-making process across the mouse brain

**DOI:** 10.1101/2023.01.01.522410

**Authors:** Elaheh Imani, Alireza Hashemi, Setayesh Radkani, Seth W. Egger, Morteza Moazami Goudarzi

**Affiliations:** Department of Computer Engineering, Ferdowsi University of Mashhad, Mashhad, Iran; The Decision Lab, Montreal, Canada; Department of Brain and Cognitive Sciences, Massachusetts Institute of Technology, Cambridge, MA 02139, USA; Department of Neurobiology, Duke University School of Medicine, Durham, NC 27710, USA

**Keywords:** intrinsic timescale, decision-making process, multiple timescales of integration

## Abstract

Across the cortical hierarchy, single neurons are characterized by differences in the extent to which they can sustain their firing rate over time (i.e., their “intrinsic timescale”). Previous studies have demonstrated that neurons in a given brain region mostly exhibit either short or long intrinsic timescales. In this study, we sought to identify populations of neurons that accumulate information over different timescales in the mouse brain and to characterize their functions in the context of a visual discrimination task. Thus, we separately examined the neural population dynamics of neurons with long or short intrinsic timescales across different brain regions. More specifically, we looked at the decoding performance of these neural populations aligned to different task variables (stimulus onset, movement). Taken together, our population-level findings support the hypothesis that long intrinsic timescale neurons encode abstract variables related to decision formation.

Furthermore, we investigated whether there was a relationship between how well a single neuron represents the animal’s choice or stimuli and their intrinsic timescale. We did not observe any significant relationship between the decoding of these task variables and a single neuron’s intrinsic timescale. In summary, our findings support the idea that the long intrinsic timescale population of neurons, which appear at different levels of the cortical hierarchy, are primarily more involved in representing the decision variable.

## Introduction

Living in natural environments entails performing a variety of tasks, each of which requires processing information over multiple timescales. For example, organisms need to extract information from visual scenes that change on the order of milliseconds and then hold and manipulate information in working memory on the order of minutes, if not hours. The heterogeneity of local microcircuits and their long-range connectivity with other circuits allow the brain to function on multiple timescales (Chaudhuri, Knoblauch, Gariel, Kennedy, & Wang, 2015). A key question is whether and how the intrinsic dynamic properties of neurons are systematically linked to their functional specializations with respect to task demands that vary in the temporal dimension.

Several studies in humans and non-human primates have demonstrated that different brain regions exhibit distinct properties in terms of the timescales of information integration. These timescales are organized hierarchically across the brain, and this hierarchical organization reflects the anatomical hierarchy of these regions (Cavanagh, Hunt, & Kennerley, 2020; Chen, Hasson, & Honey, 2015; Honey et al., 2012; Imani et al., 2022; Murray et al., 2014; Pinto, Tank, & Brody, 2022). In macaques, the timescale of intrinsic fluctuations in spiking activity has been shown to increase across the cortical hierarchy (Murray et al., 2014). A similar hierarchical organization is shown to exist across the somatosensory network (Rossi-Pool et al., 2021), and association brain areas (Gao, van den Brink, Pfeffer, & Voytek, 2020) in macaques, as well as visual areas in the mouse brain (Siegle et al., 2021). Consistent with these animal studies, studies using electrocorticographic (ECOG) signals in humans have shown that the neural timescale is consistent with the anatomical hierarchy and increases from sensory to association regions.

In addition to the overall organization of timescales across multiple brain regions that correspond to the anatomical hierarchy of those regions, there is heterogeneity in the timescale of neuronal dynamics within each functionally relevant brain region. Recent studies have revealed that the intrinsic timescale of neurons at rest, within each brain region, is associated with their response selectivity during the task (Cavanagh et al., 2020; Cavanagh, Wallis, Kennerley, & Hunt, 2016; Fascianelli, Tsujimoto, Marcos, & Genovesio, 2019; Fontanier, Sarazin, Stoll, Delord, & Procyk, 2021).

Neurons with longer timescales show higher response selectivity for temporally extended computations such as decision making and reward integration, whereas shorter timescale neurons are implicated in sensory processing. The variability of intrinsic timescales within the prefrontal cortex (PFC) in nonhuman primates predicts the involvement of these neurons in decision making, with longer timescale neurons exhibiting stronger correlations with the animal’s choice (Cavanagh et al., 2016). Populations of longer time-scale neurons were also found to represent working memory in ventrolateral PFC (Cavanagh, Towers, Wallis, Hunt, & Kennerley, 2018). Finally, longer timescale neurons in dorsolateral PFC show the stronger and more prolonged encoding of decision-related information (Fascianelli et al., 2019). Altogether, these findings suggest a potential connection between intrinsic temporal properties of neurons and their functional specialization, both across and within different brain regions.

However, the investigation of the organizational principles of brain computation across the whole brain has been limited due to the lack of physiological data with high spatiotemporal resolution from multiple brain regions. Indeed, the organizational hierarchy of neural timescales and their functional relevance has been studied only in a limited set of brain regions. Moreover, the link between the temporal properties of neurons and their functional specialization has been studied for a limited set of cognitive processes. For instance, the majority of existing work on intrinsic timescales did not investigate the temporal properties of neurons involved in perceptual decision-making.

Here, we address these gaps in the literature by investigating how the intrinsic timescale of baseline neural activity relates to the functional specialization (i.e., stimulus and choice selectivity in a perceptual decision-making task) of both single neurons and populations of neurons across the mouse brain. For this, we use two large-scale datasets recorded by Neuropixels and widefield Calcium imaging in the same visual discrimination task (Steinmetz, Zatka-Haas, Carandini, & Harris, 2019; Zatka-Haas, Steinmetz, Carandini, & Harris, 2021). Using these datasets allows us further to examine the consistency of the two recording modalities. Our population-level findings support the hypothesis that neurons with longer intrinsic timescales within and across different brain regions are more strongly implicated in the representation of the decision and stimulus.

## Results

Our analyses were performed on the activity of neurons within and across multiple brain regions of 10 mice performing a visual discrimination task (Steinmetz et al., 2019) (see Methods). Visual stimuli of varying contrast could appear on the left, right, both, or neither side of the screen on a given trial. Mice earned a water reward by turning a wheel and indicating which side had the greatest contrast or by not turning the wheel if no stimulus was presented (Figure 1 - a).

**Figure 1.**
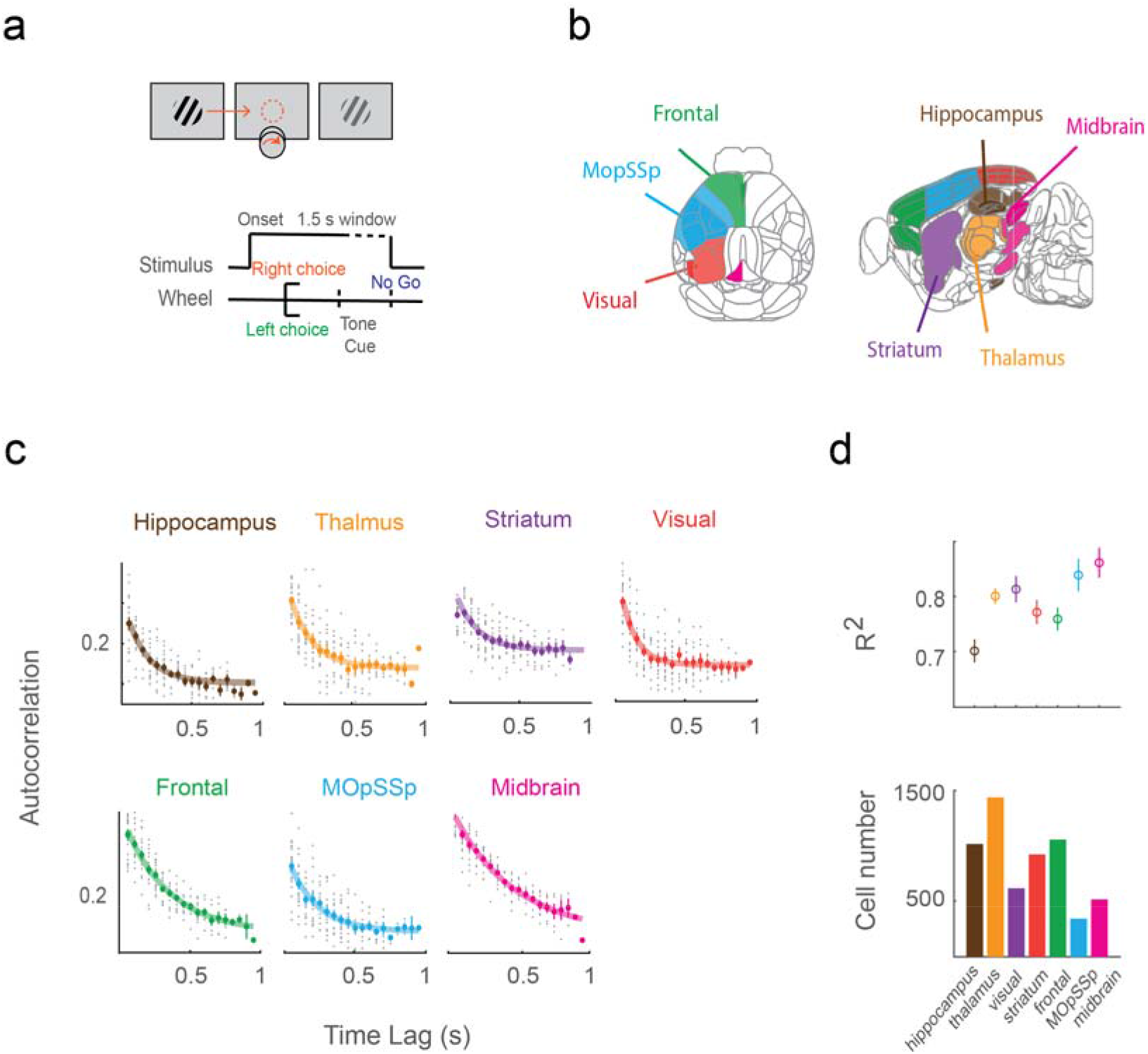
Large scale neural recording during the task and the spike count autocorrelations of the neurons within the baseline period was used to estimate the neural timescale. a) Task setup. Mice earned water rewards by turning a wheel to indicate which of two visual gratings had higher contrast or by not turning the wheel if no stimulus was presented, adapted from (*Steinmetz et al., 2019*). b) Schematic of brain regions. Recordings were made from each of the seven-colored brain regions adapted from (*Steinmetz et al., 2019*). c) Autocorrelation functions for a representative neuron from each brain region which was used to measure intrinsic timescales. d) The first row is the variance explained (R-squared) by the exponential fit to each neuron, averaged across neurons in each brain region. Error bars indicate confidence interval (CI). The second row is the number of neurons that survive all pre-selection criteria per region.

While mice performed this task, Neuropixel probes were used to collect data from approximately 30,000 neurons in 42 brain regions (Figure 1 - b & Table 1, see Methods for selection criteria). We also analyzed widefield calcium imaging data that measured the activity of the dorsal cortical regions during the same task but recorded in a separate experiment (Zatka-Haas et al., 2021). Electrophysiological and calcium imaging data formed the basis for all of our analyses.

**Table 1.** Brain regions within each group of areas in the physiological dataset according to the Allen CCF.

| **Group name** | **Regions within each group** |
| --- | --- |
| Hippocampus | POST, SUB, DG, CA1, CA3 |
| Thalamus | LP, LD, RT, MD, MG, LGd, VPM, VPL, PO, POL |
| Visual | VISp, VISrl, VISam, VISpm, VISl, VISa |
| Striatum | CP, GPe, ACB, LS |
| Frontal | MOs, ACA, PL, ILA, ORB |
| MopSSp | MOp, SSp |
| Midbrain | MRN, SCm, SCs, APN, PAG, SNr |

The “intrinsic timescale” of a neuron is defined as the decay time constant of the autocorrelation structure for that neuron. We computed this metric over a 1s baseline period (Murray et al., 2014) (see Methods). Previously, most studies of the functional significance of intrinsic timescales assigned a single timescale to an entire brain region. In contrast, we were more interested in the diversity of timescales within regions, and whether neurons with different intrinsic timescales (within particular brain regions) served distinct functional roles.

This interest stems from the fact that, in cognitive processes such as perceptual decision-making and memory retrieval, the intrinsic timescale of the neurons has been shown to partially determine their functional role (Cavanagh et al., 2018; Cavanagh et al., 2016; Fascianelli et al., 2019). However, the relationship between an individual neuron’s encoding properties and its intrinsic timescale within the brain region is poorly understood and can be broadly assessed in two ways.

First, we can determine how well two pseudo-populations of neurons with different intrinsic timescales encode a particular task variable, such as the animal’s choice or the stimuli it was exposed to (see Methods). It is worth noting that this distinction does not imply that a single neuron with a short or long intrinsic timescale encodes the stimulus or decision variable independently, but rather that populations of neurons with different intrinsic timescales contribute to the representation of a task variable to varying degrees.

Alternatively, each individual neuron may be responsible for encoding the animal’s choice or stimulus on its own. In such a case, we would expect a strong correlation between the intrinsic timescale of individual neurons within brain regions and how accurately they represent the stimulus or the animal’s choice. This paper will thoroughly examine both of these possibilities, beginning with the first.

### The heterogeneity of intrinsic timescales across brain regions

We restrict all of our subsequent analyses in the physiological dataset to neurons within the seven broad brain regions defined in Table 1. Rather than pooling the autocorrelograms of all neurons within these seven regions (which is the most common procedure), we first determined the intrinsic timescale for each neuron separately (Figure 1 - c). Despite the fact that this resulted in a more noisy fitting procedure because some neurons were inevitably poorly represented by a simple exponential decay function (Figure 1 - d, top panel), a significant proportion (~15% per area) of neurons showed a decay in autocorrelation structure that could be reliably quantified using an exponential function and were therefore included in subsequent analyses (Figure 1 - d, bottom panel, See Methods).

In the widefield calcium imaging data, we also grouped the areas in the dorsal cortex into three regions (Table 2). To be consistent with the physiological dataset, we restricted our analysis to the left hemisphere of the calcium imaging data. We then computed the timescale of the pixels by fitting the autocorrelation of the neural activity by an exponential decay function (see Methods). Similar to the physiological data, we excluded the poorly fitting pixels from further analysis (Supplemental Figure 4- a).

**Table 2.** Brain regions within each group in the calcium imaging dataset according to the Allen CCF.

| **Group name** | **Regions within each group** |
| --- | --- |
| Visual | VISp, VISpm, VISam, VISrl, VISal, VISpl |
| Frontal | MOs, ACA, PL |
| MopSSp | MOp, SSp |

We observed a high degree of within-region diversity in intrinsic timescales both in the physiological (Figure 2 - a) and calcium imaging data (Supplemental Figure 4- c). The presence of single neurons with a diverse range of timescales across different brain regions suggests that this observed heterogeneity has a functional relevance. This finding prompted us to examine the functional properties of neurons with different intrinsic timescales within brain regions.

**Figure 2.**
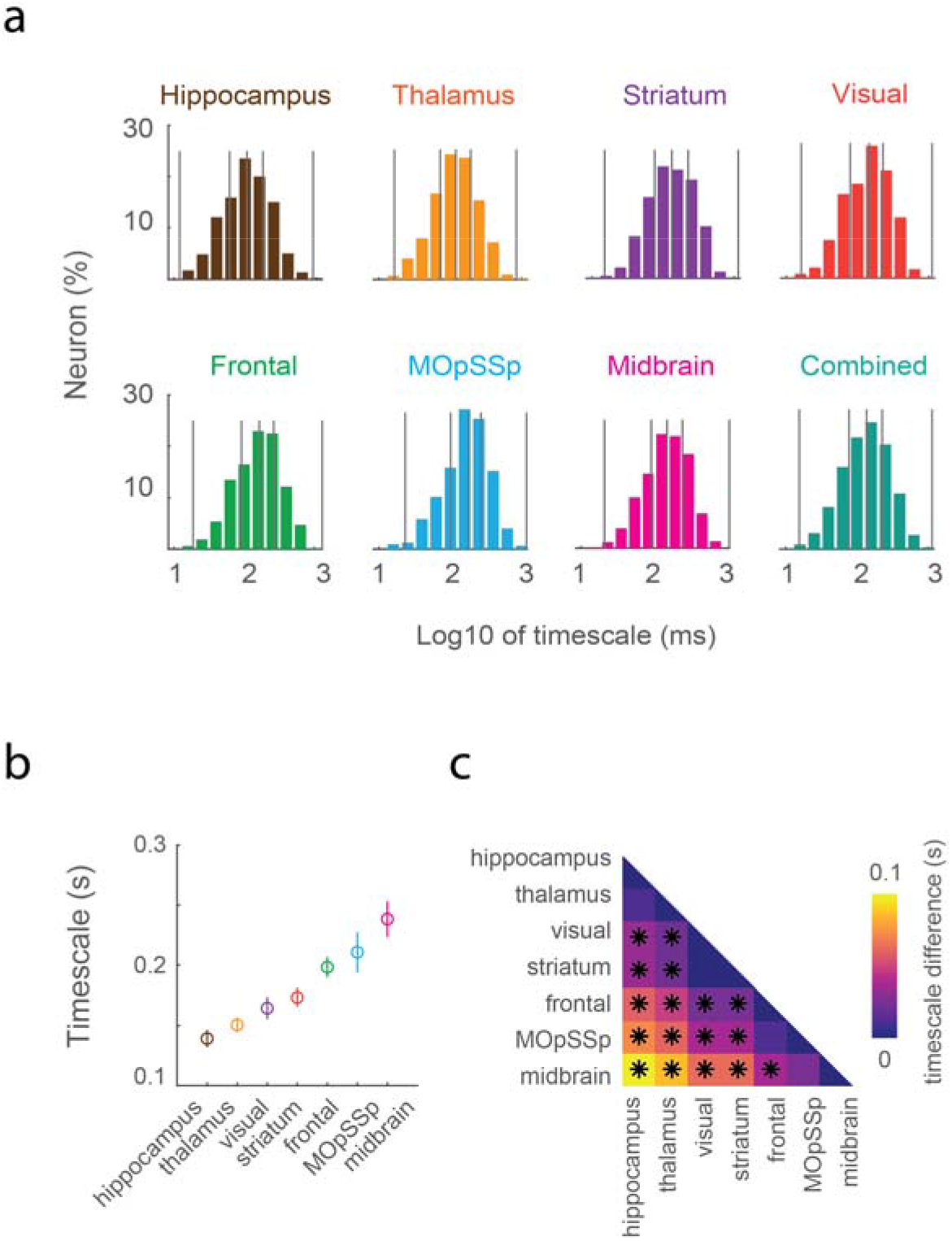
Heterogeneity of single-neuron timescales within each brain region and the hierarchical ordering of the timescales across regions. a) Distribution of intrinsic timescales for each brain region. Vertical lines represent the quartiles. b) Intrinsic timescales averaged across neurons within each brain region and sorted in an ascending manner. Errors indicate CI. c) Pairwise differences in intrinsic timescales across brain regions. Marker ‘*’ symbolizes significant differences between region’s timescale in a permutation test corrected for multiple comparisons.

As a first step, we computed the average of intrinsic timescales within each brain region (Figure 2 - b & c, Supplemental Figure 4- b). We arranged brain regions in ascending order based on their average intrinsic timescale. We will hereafter refer to this ordering as the “intrinsic timescale hierarchy”, in short, ITH. As can be observed from these figures, the timescale hierarchy of the regions in the dorsal cortex is consistent in both datasets. In the physiological data, this hierarchy begins with hippocampal neurons and ends with midbrain neurons. Moreover, it is loosely correlated with the general functions of these regions. For example, brain regions implicated in motor and decisionmaking activities had intrinsic timescales that were longer on average than that of visual neurons.

### Relationship between the Neural Intrinsic Timescale and Decision-making

Given that the integration of sensory evidence is a temporally prolonged activity, we hypothesize that single neurons with longer intrinsic timescales within and across different brain regions are more strongly implicated in the representation of the animal’s decision. As a first attempt, we addressed this hypothesis by splitting the neural population within each brain region based on its intrinsic timescale and performing a population decoding analysis of both the stimulus and the animal’s choice for each of these splits.

The direction of wheel movement during correct trials was the first task variable we decoded. Across the ITH, we observed a broadly improved representation of this measure (Figure 3 - a & b). In other words, brain regions that had a higher average intrinsic timescale represented the animal’s movement more accurately than those with a lower average intrinsic timescale (Figure 3 - b).

**Figure 3.**
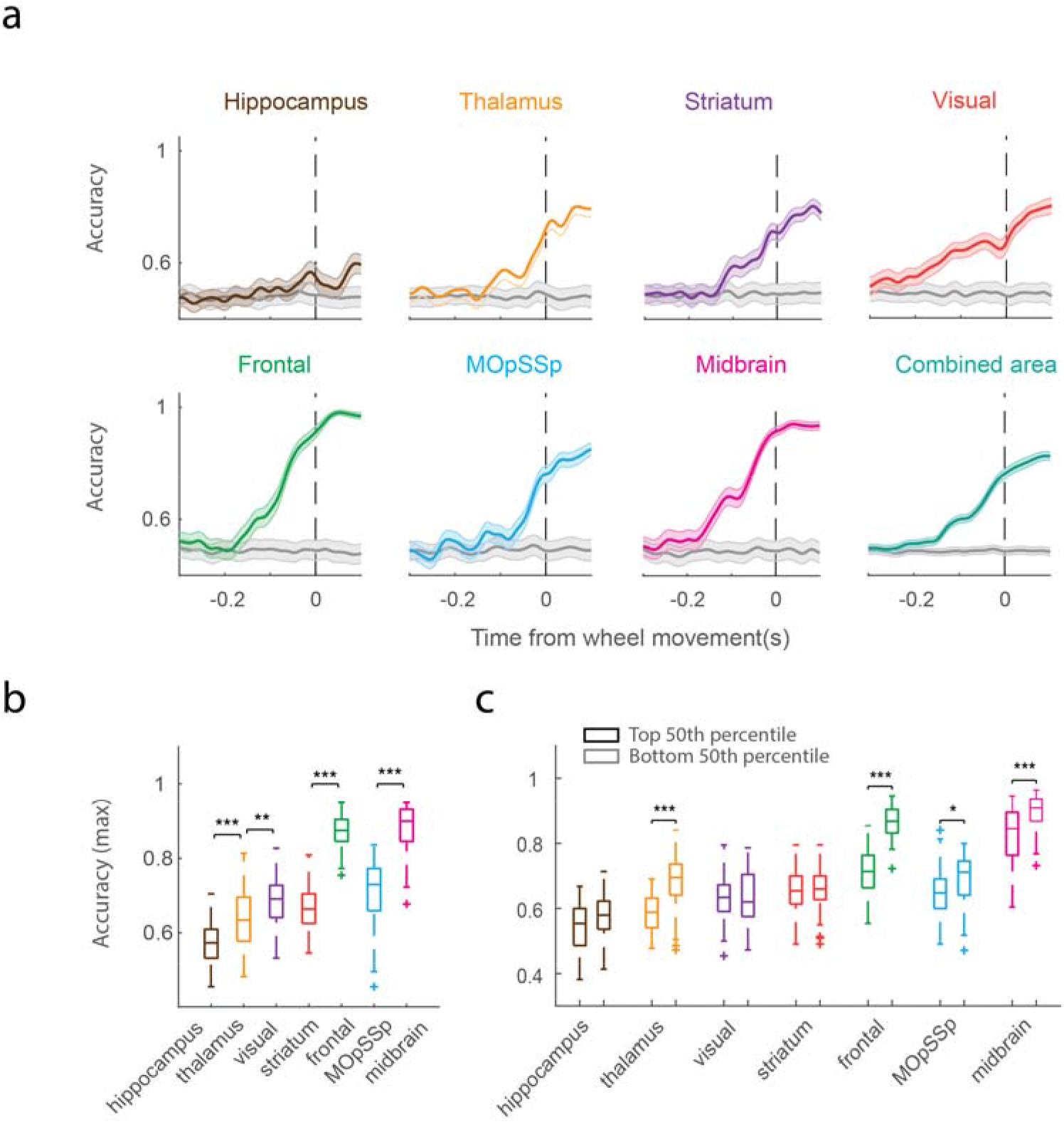
Population decoding of movement for total, short, and long-timescale neurons. a) Timecourse of population decoding of movement, aligned to the onset of the wheel turn. Shaded areas indicate CI. b) Maximum population decoding of movement in different brain regions sorted according to ITH. c) Maximum population decoding of movement in neurons with short and longtime scales (median split) sorted according to ITH. Across all areas (except for visual), longer timescale neurons (neurons that fell into the Top 50th percentile) represented information about the animal’s movement with higher accuracy than shorter timescale neurons. The signs ‘***’ and ‘*’ symbolize p-value < 0.001 and p-value < 0.05 produced by a Wilcoxon rank-sum test corrected for multiple comparisons.

Furthermore, when we separated neurons within these brain regions based on their intrinsic timescale, we found that long intrinsic timescale neurons were better at decoding wheel movement, specifically within brain regions that are heavily implicated in decision-making and learning (Figure 3 - c, e.g., striatum and frontal).

### Relationship between the Neural Intrinsic Timescale and Stimulus Decoding

We also hypothesized whether or not the neural population with a longer timescale within each brain region can predict the stimulus contrast level of the contralateral stimulus better than shorter ones. One significant limitation of this decoding measure is that it conflates choice decoding with stimulus decoding. In other words, a neuron or population of neurons may respond preferentially to the stimulus or decision variable. However, because there is a strong correlation between the strength of the stimulus and the animal’s choice, it is not easy to distinguish whether the population of neurons more strongly represents either of these variables. Therefore, we devised a method to separate a neuron or a neural population’s response to the stimulus from its response to the choice (see Methods). We will evaluate this metric in subsequent sections.

Accordingly, we divided the trials into 12 groups based on three different animal choices and four contrast levels. We then compared the activity of these neural populations between trials where the right stimulus contrast was zero, and the right stimulus contrast was non-zero within each group. The combined accuracies across 12 groups were used for computing final stimulus decoding. We found that we could decipher visual information at above-chance levels by decoding the activity of both short and long intrinsic timescale neurons in these locations (Figure 4 - a).

**Figure 4.**
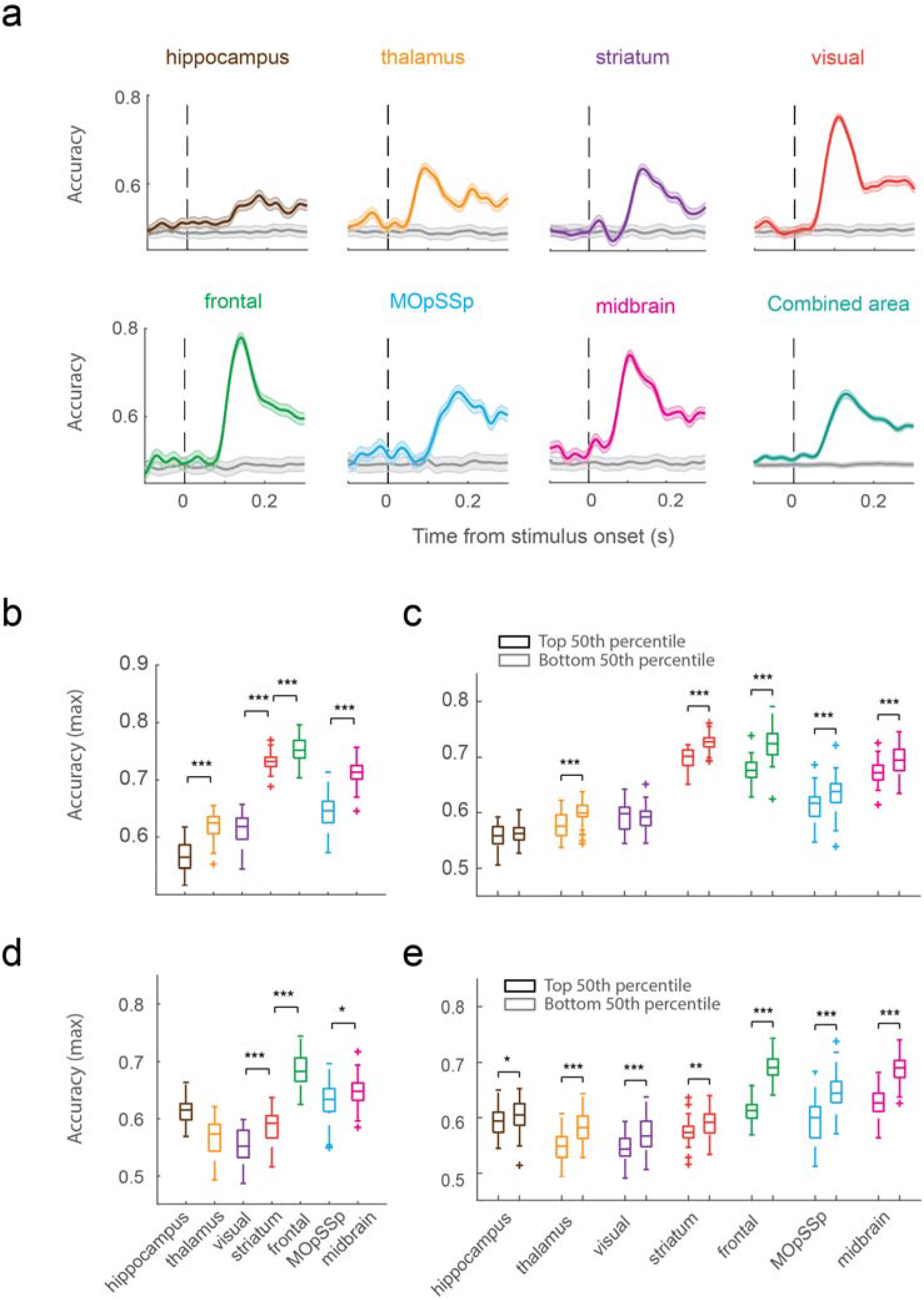
Population decoding of stimulus and decision for total, short, and long-timescale neurons. a) Timecourse of stimulus population decoding. Shaded areas indicate CI. b) Maximum decoding accuracy of the stimulus for different brain regions sorted according to ITH. c) Median split of the maximum population for stimulus decoding. d) Maximum decoding accuracy of decision across brain regions. e) Median split of the population for the decision decoding. The signs ‘***’, ‘**’ and ‘*’ symbolize p-value < 0.001, p-value < 0.01 and p-value < 0.05 in a Wilcoxon rank-sum test corrected for multiple comparisons.

Furthermore, for the majority of the brain regions we evaluated (5 out of 7), long intrinsic timescale neurons decoded the stimulus with greater accuracy than short ones (Figure 4 - c). And aside from visual areas, the midbrain and frontal areas exhibited strong stimulus decoding, which is consistent with their well-documented roles in perceptual decision-making (Coen, Sit, Wells, Carandini, & Harris, 2021; Scott et al., 2017; Steinmetz et al., 2019; L. Wang, McAlonan, Goldstein, Gerfen, & Krauzlis, 2020; Zatka-Haas et al., 2021). Consistent with these findings, our calcium imaging results also support that longer intrinsic timescale neurons contribute more to stimulus decoding across brain regions (Supplemental Figure 5- e).

### Relationship between the Neural Intrinsic Timescale and Decision (Detect Probability) Decoding

We sought additional evidence to support our hypothesis that long intrinsic timescale neurons encode abstract variables associated with decision formation. As a result, we used a metric to decode the animal’s choice independent of stimulus strength. We will refer to this decision-making metric as “population detect probability” (population DP). As mentioned before, there is a strong correlation between stimulus and animal choice. So similar to the stimulus decoding, we used the combined condition decoding approach by splitting the trials into 12 groups according to the different combinations of right and left stimulus contrast levels. We then separated the trials within each group into Hit (correctly turning the wheel) and Miss classes.

We generally expect higher-level regions to be involved in representing the correct perceptual decision by reading out noisy information from lower levels of the hierarchy to form a choice. Consistent with this, we found that decision-making and learning areas had a higher average population DP than sensory areas. Furthermore, we then divided the population into long and short intrinsic timescales; we discovered that neurons with longer intrinsic timescales (Top 50th percentile) could predict the animal’s decision more accurately than neurons with shorter intrinsic timescales (Figure 4 - e, Bottom 50th percentile). Consistent with these findings, our calcium imaging results also lend support to the idea that longer intrinsic timescale neurons contribute more to decision decoding across brain regions (Supplemental Figure 5- f).

### Intrinsic neural timescales and single neuron decoding of the animal’s choice and stimuli

Our initial hypothesis was that neurons with longer intrinsic timescales would more strongly represent the decision-making process than neurons with shorter intrinsic timescales. This follows from the fact that sensory evidence integration is a temporally prolonged activity. In the previously reported analyses, we addressed and largely confirmed this hypothesis by splitting the neural population based on its intrinsic timescale and performing a population decoding analysis. Here we present a secondary approach; namely, we also wanted to find out if single neurons with long or short intrinsic timescales represented information regarding the animal’s decision or the stimuli presented to the animal.

The first step was to determine which subset of neurons more strongly encoded the stimulus and which neurons more strongly encoded the choice (i.e., the animal’s decision to move or not move the wheel). This was a significant challenge because, during the course of a given task, individual neurons in the cortex may encode a variety of distinct task variables.

This simultaneous representation of diverse components of a task is referred to as a ‘mixed’ representation, and it is difficult to decipher. Kobak, Brendel et al. (2016) developed a data analysis tool called demixed PCA (dPCA) to address this problem (Kobak et al., 2016). Similar to other dimensionality reduction approaches, dPCA breaks down the activity of a population of neurons into a limited number of components. Unlike other approaches, each dPCA component relates only to a particular task-relevant variable, making it easy to interpret. For example, the component of the neural population representing the animal’s choice can be differentiated from the one that carries information about the stimuli it was exposed (Figure 5 – a & Supplemental Figure 1).

**Figure 5.**
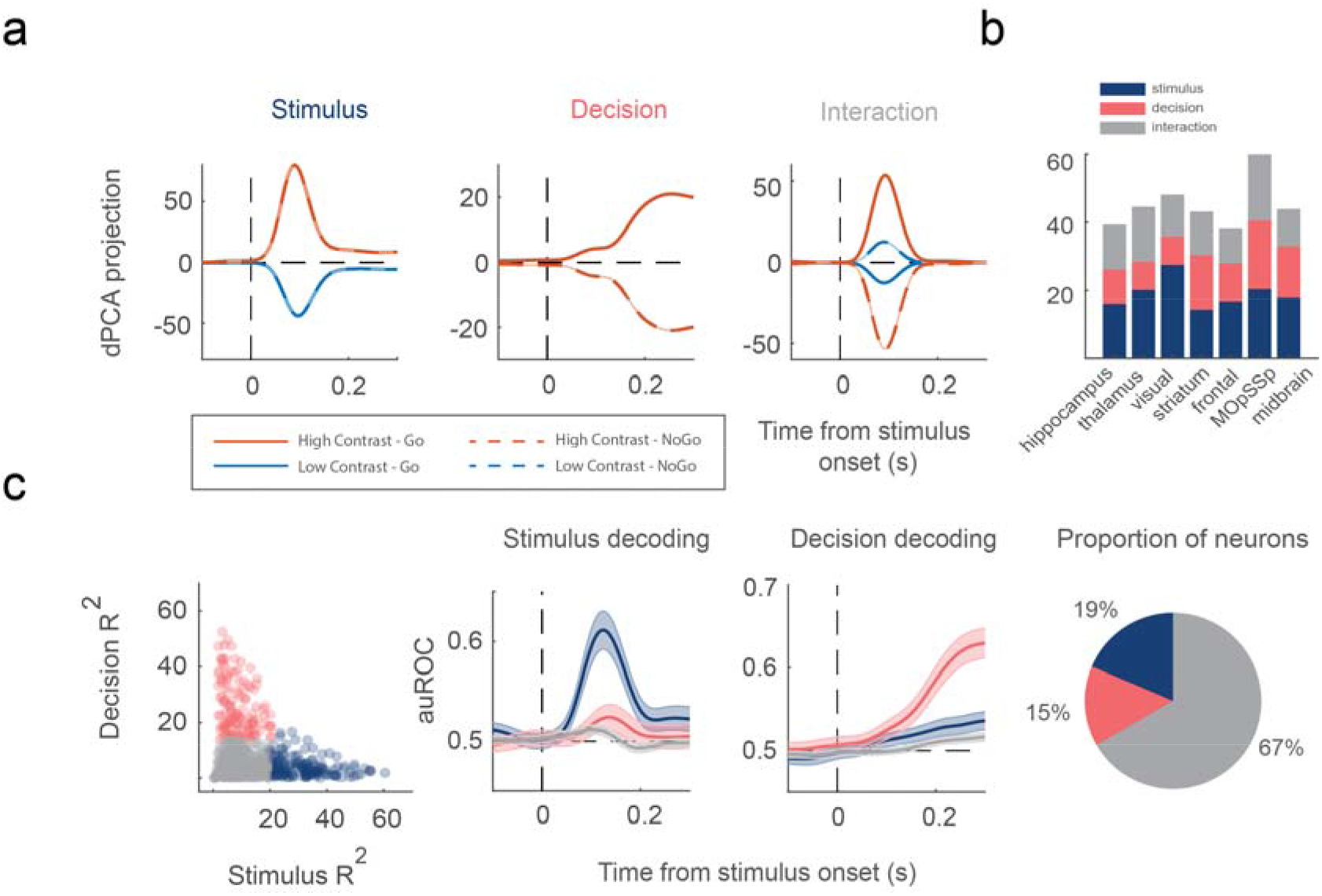
Demixing neural population into task-related components. a) Each subplot shows a demixed principal component (corresponding to the stimulus, the decision, and interaction). Each subplot has four lines corresponding to four conditions (see legend). b) Explained variance of the individual demixed principal components for each brain region. c) Clustering of three neuron subtypes: stimulus (blue), decision (pink), and other (grey) based on demixed principal components (left). auROC validation of the three neuron subtypes (right). Shaded areas indicate CI. Stimulus neurons (blue) decode the stimulus much better than the other neuron subtypes. Decision neurons (pink) encode the decision (or “detect probability) better than the other neuron subtypes. d) Proportion of different subtypes for this example region (visual).

We clustered neurons using the stimulus and choice components of dPCA (see Methods) based on whether they responded more strongly to the stimulus or the choice. Accordingly, the activity of the neurons was projected to the stimulus and choice components using the decoder matrices of dPCA. We then reconstructed the neural activity using the choice and stimulus encoder matrices. Using the neurons’ explained variance (R-squared) and fuzzy C-Means algorithm, we indicated whether they responded more strongly to the stimulus or the choice. We discovered that neurons across brain regions generally fell into one of three clusters: those best represented by the stimulus component, the choice component, or their interaction. As a result, we used clustering to determine whether a neuron was stimulus-selective, choice-selective, or neither (Figure 5 – b & Supplemental Figure 2 - a).

The measures we used to evaluate single neuron decoding were based on an auROC metric (Figure 5 - c & Supplemental Figure 2 - b & c). This method is commonly used to calculate the difference between spike count distributions across trials (see Methods). It can be used to calculate a neuron’s sensitivity to parametrically varying stimuli (stim decoding) or to quantify the relationship between a neuron’s activity and the animal’s choice (detect probability, DP). Due to differences in firing rate associated with each stimulus strength, DP was independently calculated for each stimulus value.

Previous research has found that DPs typically increase in value as one approaches the site of decision making and that DP values in early sensory areas are significantly lower than those in sensory areas. Given that intrinsic timescales vary according to brain region, we investigated whether there was also a relationship between these auROC-based measures of choice and stimulus selectivity and individual neuron intrinsic timescales.

We discovered that a significant proportion of the neurons we examined were either stimulus or choice selective. The proportion of each type of neuron within each brain region varied greatly (Figure 6 - a). We hypothesized that individual neurons with longer timescales ought to represent decision variables. Put another way; there should be a positive relationship between neural timescales and single neuron decoding of the animal’s decision (detect probability). However, no statistically significant relationship was discovered between either the decision or stimulus decoding properties of individual neurons (within brain regions) and their intrinsic timescale (Figure 6 - a, middle and right panel, Table 3). Moreover, we did not observe a significant difference between how short and long intrinsic timescale neurons represented the stimulus or choice signal at any point after stimulus onset (Supplemental Figure 3). Furthermore, we discovered that there was no evidence of a stimulus or decision decoding hierarchy across the brain regions we studied either (Figure 6 - b & c).

**Figure 6.**
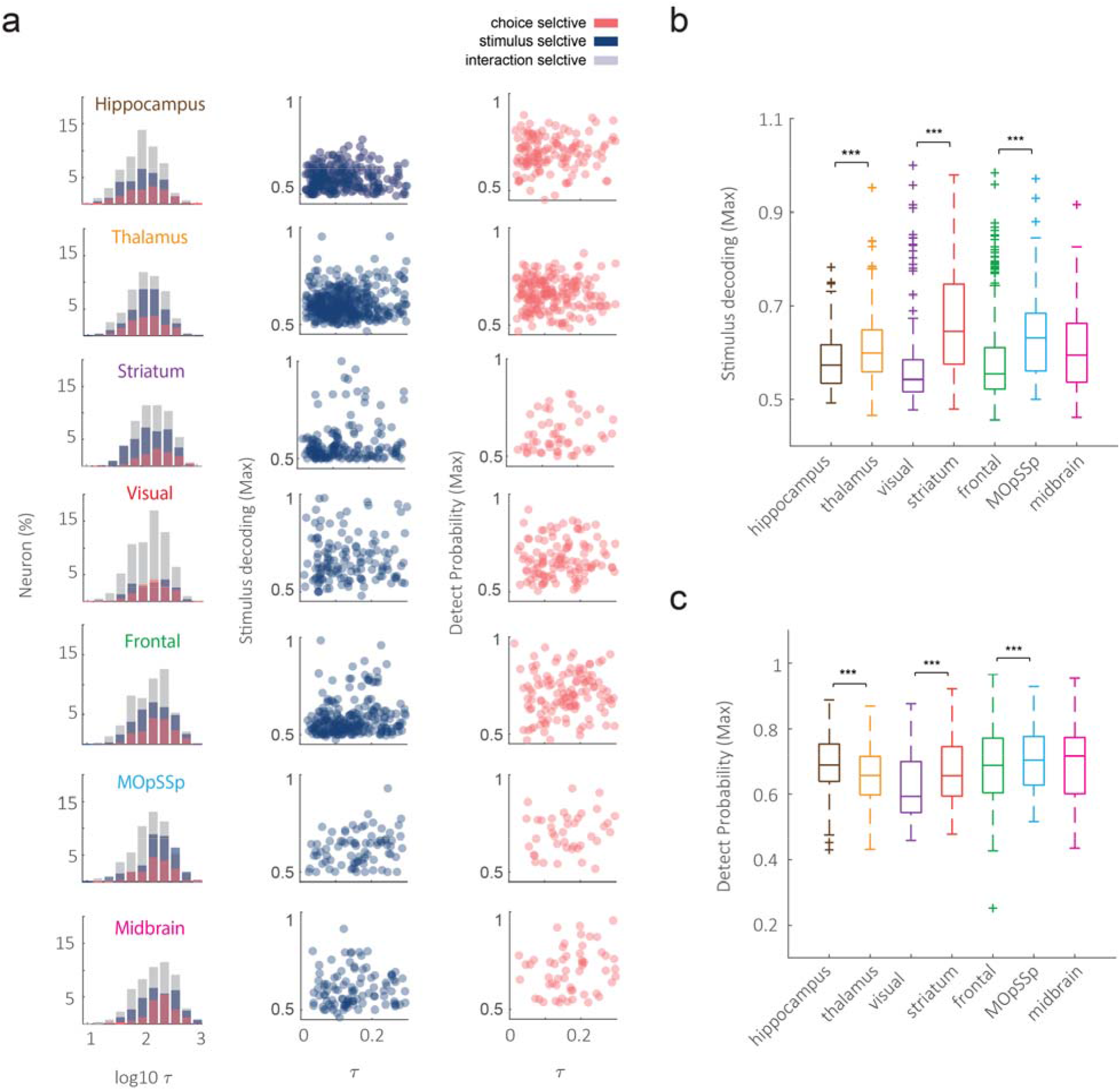
Decoding accuracy of stimulus and choice selective neurons. a) the first column indicates the distribution of timescale for different types of neurons. The second and third columns demonstrate the scatter plots for individual neuron stimulus decoding and DP, respectively. b) Maximum stimulus decoding for different brain regions sorted according to ITH. c) Detect probability for different brain regions according to the ITH ordering. The signs ‘***’ and ‘**’ symbolize p-value < 0.001 and p-value < 0.01 produced by the Wilcoxon rank-sum test corrected for multiple comparisons.

**Table 3.**
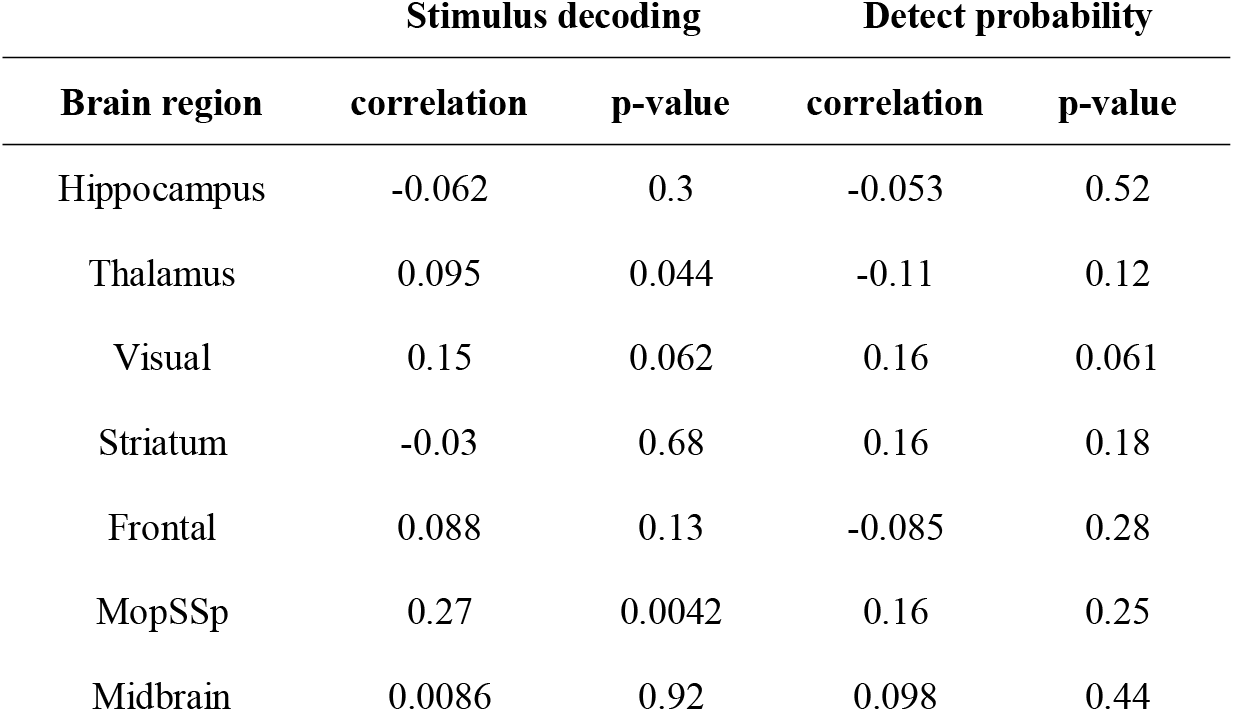
Pearson’s correlation between intrinsic timescale and measures of single neuron decoding.

In conclusion, the relationship between neural timescales and their role in decision-making was observed at the population level but not at the single neuron level. This relationship was especially strong in brain regions involved in decision-making and learning. Neurons in these brain regions play a critical role in dynamically tracking value and volatility in the environment (Massi, Donahue, & Lee, 2018).

## Discussion

We have demonstrated that intrinsic timescales have high predictive power when it comes to characterizing the decision-making and stimulus encoding properties of populations of neurons. The brain regions we examined contained single neurons with a wide range of timescales. Nonetheless, these brain regions varied in their average intrinsic timescales, giving rise to a hierarchical organization based on temporal dynamics. However, we did not find strong evidence for a systematic relationship between individual neurons’ intrinsic timescales and their functional specialization.

Within brain areas, we also identified functional variations between neurons with varying intrinsic timescales. We found that neurons with more sustained activity (longer intrinsic timescales) were more likely to be involved in perceptual decision-making at the population level but not at the single neuron level. Taken together, our findings at the population level lend support to computational theories that attribute evidence integration to highly recurrent attractor networks (X.-J. Wang, 2002).

There are two potential biophysical mechanisms that could give rise to longer intrinsic timescales (Murray et al., 2014). One potential mechanism is stronger synaptic connections mediating recurrent excitement, which slow intrinsic dynamics by partially negating leak (Goldman, Compte, & Wang, 2010). Another mechanism is an increase in the number and density of excitatory synapses onto pyramidal cells across the cortical hierarchy (Elston, 2003). Modeling studies have revealed that strong recurrent connections enable neurons to exhibit persistent activity in working memory (Kim & Sejnowski, 2021; Zylberberg & Strowbridge, 2017), as well as the ability to accumulate information slowly during decision-making (X.-J. Wang, 2002).

A hierarchy of timescales may offer a distinct computational advantage for the mammalian nervous system. Previous studies have shown that when basic recurrent neural networks (RNNs) are task-optimized, a hierarchy of neural timelines (within brain regions) automatically emerges. In particular, RNNs with a hierarchy of timescales have been shown to be effective at learning tasks that are a re-composition of previously learned sub-goals (Han, Doya, & Tani, 2020).

There is one important caveat with our study: while decoding can reveal how much information a neuronal population has about a function, high decoding accuracy does not necessarily imply that a brain area is directly involved in processing that function. Furthermore, while our calcium imaging results lend support to the idea that longer intrinsic timescale neurons contribute more to sensory processing and decision-making across brain regions, we did not observe a similar functional hierarchy across brain areas to what we found in single units. This disparity could be explained by the fact that Neuropixels record neural activity from multiple layers of the cortex, whereas calcium imaging only records neural activity from the first layer of the cortex. This discrepancy may possibly be explained by the fact that the number of neurons included in MOpSSp after preprocessing was much smaller than the number of calcium imaging channels.

To summarize, in this paper, we identified a functionally relevant heterogeneity in intrinsic timescales, which enables the brain to perform tasks that necessitate longer-term representations of choice and stimulus information. We found that this result holds true across brain regions and modalities. We made several novel contributions to the existing body of work. We devised a method to decompose the contribution of intrinsic timescales to decision variables from their representation of the stimulus. The results presented in this publication are more reliable than prior research on intrinsic timescales, which did not always adequately separate stimulus and choice contributions to population brain activity.

## Materials and Methods

### Behavioral Task

We analyzed a dataset made freely available by (Steinmetz et al., 2019). This dataset contains behavioral and physiological data collected from ten mice over the course of 39 sessions on a visual discrimination task (two-alternative unforced choices). Mice rested on a plastic apparatus with their forepaws on a rotating wheel, surrounded by three computer monitors. To begin the trial, the animal briefly held the wheel. Then, using 16 various conditions, visual stimuli (Gabor patch with sigma 9 and 45° direction) with four grading levels might be exhibited on the right, left, both, or neither screen.

The animal did not need to move its head to perceive the stimuli because they were presented in the mouse’s central monocular zones. Mice were rewarded by moving the wheel such that the stimulus with the highest contrast moved to the center of the screen or by not turning the wheel at all if no discernable stimuli were shown. Otherwise, they received a white noise signal to indicate an improper wheel movement. So, three types of outcomes (turn right, turn left, no turn) led to the reward according to the stimulus presentation.

After the stimulus presentation, there was a random delay interval of 0.5–1.2 s during which the mouse could freely turn the wheel, but visual stimuli were locked in place, and incentives were unavailable. At the conclusion of the delay interval, an auditory tone cue (8 kHz pure tone for 0.2 s) was supplied, at which point the visual stimulus position became coupled to wheel movement.

## Electrophysiological data

### Recordings

Neuropixel electrode arrays were utilized to record approximately 30,000 neurons in 42 in the left hemisphere brain areas during the task. Due to the capacity of a single probe to record from numerous brain regions and the usage of many probes concurrently, each session produced data recorded from many areas simultaneously. We divided the regions into seven main groups according to the Allen Mouse Brain Common Coordinate Framework (CCF) atlas (Q. Wang et al., 2020) (Figure 1 - b). All the analyses were performed on these groups of regions.

### Data preprocessing

We excluded cells from our decoding analysis based on the neuron’s timescale goodness of fit. Spikes were binned at 0.005s and smoothed using a half-gaussian kernel with a standard deviation of 0.02s. Across all trials and time points, data from all neurons were z-scored by subtracting the mean and dividing by the standard deviation calculated during the baseline period (−0.9s to −0.1s, stimulus aligned).

All analyses were carried out using customized MATLAB code. Statistical analyses, SVM classification, and fuzzy C-means clustering were performed using MATLAB toolboxes. We also used the open-source dPCA toolbox (Kobak et al., 2016) to decompose neural activity into different task-related variables.

### Intrinsic timescale

Our measure of intrinsic timescale was based on the spike count autocorrelation function proposed by (Murray et al., 2014). The spike count of each neuron was first binned at *Δ* = 0.05s using nonoverlapping windows during a 1s baseline period over Go trials. Next, the across trial Pearson’s correlation was computed between each pair of time bins *iΔ* and *jΔ* (*j > i*) separated by time lag (*j – i*)Δ. The autocorrelation values follow an exponential decay from lower to the higher time lags. So, the intrinsic timescales were estimated by fitting an exponential decay function to the autocorrelation values of each neuron, as follows:

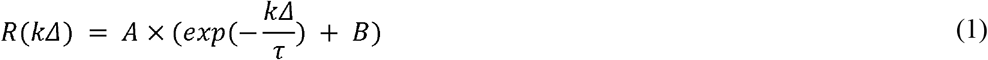

Where *kΔ* is the time lag, *τ* is the intrinsic timescale, A is the amplitude, and B denotes the contribution of timescales longer than the observation window. Some neurons had low correlation values at shorter time lags which may reflect negative adaptation (Murray et al., 2014). Fitting began with the lag with the greatest autocorrelation reduction to overcome this feature. Using five different initial parameter values, the Levenberg-Marquardt method was used to fit equation (1) to the autocorrelation values. The model with the lowest mean square value (MSE) was chosen as the best model for describing the decay of a particular neuron’s autocorrelation function.

The temporal autocorrelation of some neurons could not be described by an exponential decay. Therefore, we automatically excluded neurons that can not satisfy the following criterias: (1): minimum average firing rate of 1HZ during 1s baseline period across Go trials, (2): minimum R-squared value of 0.4 for the exponential decay fitting, (3): 0.01s < *τ* < Is. Accordingly, about half of the neurons satisfied these requirements and were further analyzed by the visual inspection process. During this phase, neurons were visually inspected and eliminated if their temporal autocorrelation function was not well-captured by an exponential decay function. A significant proportion of neurons (~15% per region) passed both the automated and visual inspection procedure and were thus selected for further analyses.

### Population Decoding

We used a binary support vector machine (SVM) classifier with a linear kernel function to decode task variables from neural activity. We measured how neural activity predicted the direction of wheel movement, whether or not the animal turned the wheel, and the stimulus that the animal was exposed to using this classifier.

### Decoding the direction of wheel movement

In order to decode the direction of wheel movement, neurons within each brain region were combined to form a pseudo-population by randomly selecting correct trials for each direction (without replacement, sample size = 10). Sessions with less than ten correct trials for each direction were excluded from subsequent decoding analyses. We measured the accuracy of the classifier using a 3-fold cross-validation procedure over 50 iterations of trial sub-sampling at each time point. We used the activity of neurons in a window ranging from −0.3 s to 0.1s (aligned with wheel movement) for this analysis.

### Decoding the contralateral stimulus

By design, the encoding of the stimulus is highly correlated with the animal’s choice. As a result, the decoding of wheel movement direction reflects an interaction between both the stimulus and choice variables. To address this limitation, we performed our decoding analyses within different choices and stimulus conditions and combined their accuracies. Because the data was recorded from the left hemisphere, we only decoded the stimulus on the right screen. The trials for contralateral stimulus decoding were divided into 12 groups based on the animal’s choice alternatives (right, left, NoGo) and four left stimulus contrast levels (100, 50, 25, 0) (Zatka-Haas et al., 2021). We used the activity of neurons in a window ranging from −0.1s to 0.3 s (aligned with stimulus onset) for this analysis.

Following that, the trials within each group were divided into two categories: (1) trials with a right stimulus contrast level of zero; (2) trials with a right stimulus contrast level greater than zero. We then generated a pseudo-population for each condition by selecting ten trials from each class at random across sessions. If a session did not meet the trial cutoff criteria for a specific condition, it was removed from the pseudo-population for that condition. We used a 3-fold cross-validation procedure to assess the performance of 12 classifiers at each time point. The final accuracy for each time point was then calculated by averaging the accuracies of these 12 classifiers. To compute confidence intervals, this sampling procedure was repeated 50 times.

### Detect probability

We next measured how the neural activity encodes whether or not the animal turned the wheel correctly and referred to this measurement as ‘Detect probability’ (Hashemi, Golzar, Smith, & Cook, 2018). We similarly divided the trials into 12 groups according to the different combinations of right and left stimulus contrast levels (0, 25, 50, 100), ignoring equal contrast pairs. The trials within each group were separated into Hit (correctly turning the wheel) and Miss classes. We then sub-sampled the trials within each class randomly without replacement with sample size five to prepare 12 pseudo-populations across sessions. The sessions with a trial number lower than five within each condition were excluded from the condition’s pseudo-population. We then measured the performance of the decoder by combining the accuracy of 12 classifiers. Classifier accuracy was assessed using a 3-fold cross-Validation procedure applied to each time point during the stimulus epoch (−0.1s to 0.3s stimulus alignment). The sampling process was repeated 50 times to measure the classifier’s confidence interval.

We also measured how the neural activity encodes task variables as a function of the neuron’s intrinsic timescale. Accordingly, we split the neurons within each brain region into short and long timescales according to their median timescale values. For each group of neurons, we repeated the above procedures for measuring the performance of each decoder.

### Single neuron decoding

The single neuron decoding analysis was performed using the area under the receiver operating characteristic (auROC) analysis (Britten, Newsome, Shadlen, Celebrini, & Movshon, 1996). This method is commonly used to calculate the difference between spike count distributions across two conditions. As mentioned before, the encoding of the stimulus and the animal’s choice are highly correlated. Therefore, similar to the population decoding, we utilized the combined condition auROC analysis to resolve this limitation for decoding the stimulus and detect probability during the stimulus epoch (−0.1s to 0.3 s, aligned to stimulus onset).

We only decoded the contralateral stimulus by dividing the trials into 12 groups according to the three animal choice alternatives and four left stimulus contrast levels. We then measured the neuron’s stimulus selectivity by applying the auROC analysis per condition. The final neuron’s stimulus selectivity was measured by the weighted average across 12 auROC values per time point.

We also assessed how a single neuron could encode whether or not the animal’s turned the wheel (Detect probability). Similar to the population decoding, we first divided the trials into 12 groups based on different combinations of right and left stimulus levels ignoring the equal contrast pairs. We then evaluated the neuron’s detect probability by taking the weighted average of auROC analysis across conditions.

### Task-related neural data decomposition

Most neurons encode different types of task information and exhibit a mixed selectivity. The complexity of single-neuron responses can conceal the type of information expressed by them and how it is represented. To circumvent this constraint, we used demixed principal component analysis (dPCA) (Kobak et al., 2016). Briefly, dPCA decomposes the population neural activity into a few latent components, with each capturing a specific aspect of the task. The compressed subspace derived from dPCA captures the bulk of the variation in the data while also decoupling different task-related components, such that each component captures variance primarily associated with a single task variable.

We first prepared a matrix containing the marginalized population’s neural activity for different stimulus and choice conditions to construct the latent subspace. Accordingly, we divided the trials into four groups based on the contralateral contrast levels. Within each contrast level, we computed the average firing rate of neurons based on whether or not the animal turned the wheel. The neural population matrix *X*_*N*×4:×2×*T*_ contains the activity of N neurons over four different stimulus contrast levels and two alternative animal wheel movement statuses with T time points during the stimulus epoch (−0.1s to 0.3s). We then applied dPCA to the neural population to construct a latent subspace containing 20 components characterizing the stimulus (*X_st_*), decision (*X_dt_*), stimulus-decision interaction (*X_sdt_*), and condition-independent information (*X_t_*) by minimizing the following loss function (Kobak et al., 2016):

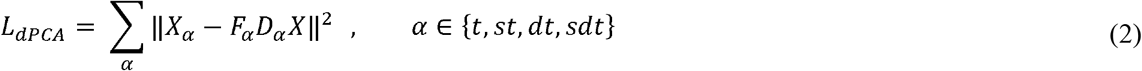

Where F and D refer to the decoder and encoder for each task variable *α*. Using the estimated decoder and encoder for each task variable, we can compute the explained variance (R-squared) of the neurons using the following equations:

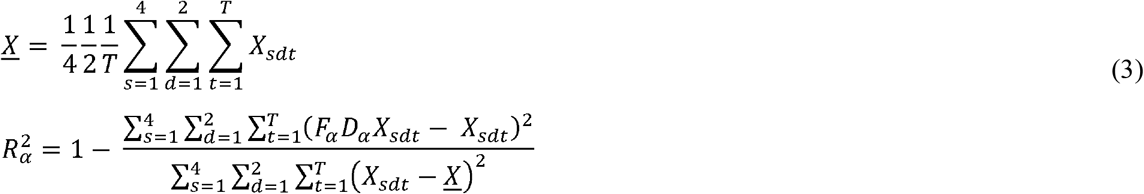

Where 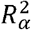 is the explained variance of each task variable *α* ∈ {*t,st,dt,sdt*}. The neurons within each brain area were then clustered (number of clusters = 3) based on their stimulus and choice-related R-squared values 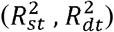, using the fuzzy C-means clustering algorithm (Bezdek, 2013).

## Widefield calcium imaging data

### Image acquisition and preprocessing

We analyzed the calcium image data made public by (Zatka-Haas et al., 2021) with the same task protocol and animal subjects as the electrophysiological dataset. Details on the data acquisition and preprocessing steps are described in (Zatka-Haas et al., 2021). The widefield calcium images were acquired through a fluorescence macroscope (Scimedia THT-FLSP) containing an sCMOS camera (PCO Edge 5.5) with a frame rate of 70HZ and pixel size 21.7μm. The authors mapped each mouse’s average cortical fluorescence activity to the Allen Common Coordinate Framework (CCF) using the grid centered on bregma. They also applied different preprocessing steps for denoising and normalizing the images.

We created a grid box at the CCF space covering the whole dorsal cortex with a grid size of 10μm. We then mapped the mouse’s image to this grid box by spatially binning pixels falling within each grid. Across all trials and time points, the activity of pixels was z-scored by subtracting the mean and dividing by the standard deviation calculated during the baseline period (−1s to 0s, stimulus aligned).

### Timescale analysis

We measured the intrinsic timescale using the autocorrelation structure of fluorescence activity during the baseline period. The activity of each pixel was linearly interpolated at 20 time-bins separated by 0.05s time lag during a 1s baseline period over Go trials. Next, the across trial Pearson’s correlation was computed between each pair of time bins *iΔ* and *jΔ* (*j > i*) separated by time lag (*jΔ - iΔ*). The autocorrelation values follow an exponential decay from lower to higher time lags. So, similar to the physiological data, the intrinsic timescales were estimated by fitting an exponential decay function to the autocorrelation values of each pixel.

The temporal autocorrelation of some pixels could not be described by an exponential decay. Therefore, we automatically excluded pixels having a timescale lower than 0.01s or higher than 1s. The remaining pixels were further analyzed by the visual inspection process. During this phase, pixels were visually inspected and eliminated if their temporal autocorrelation function was not well-captured by an exponential decay function. A significant proportion of pixels (~50%) passed both the automated and visual inspection procedure and were thus selected for further analyses.

### Decoding analysis

Similar to the single neuron decoding analysis, we used combined condition auROC analysis to decode the stimulus and detect the probability of each pixel during the stimulus epoch (−0.1s to 0.3s, aligned to stimulus onset). Consistent with the previous analysis, we only considered pixels at the left hemisphere. We performed the auROC analysis per session, and final decoding accuracy was computed by taking the average of the auROC values of each pixel and time point across sessions

## Supplemental Figures

**Supplemental Figure 1.**
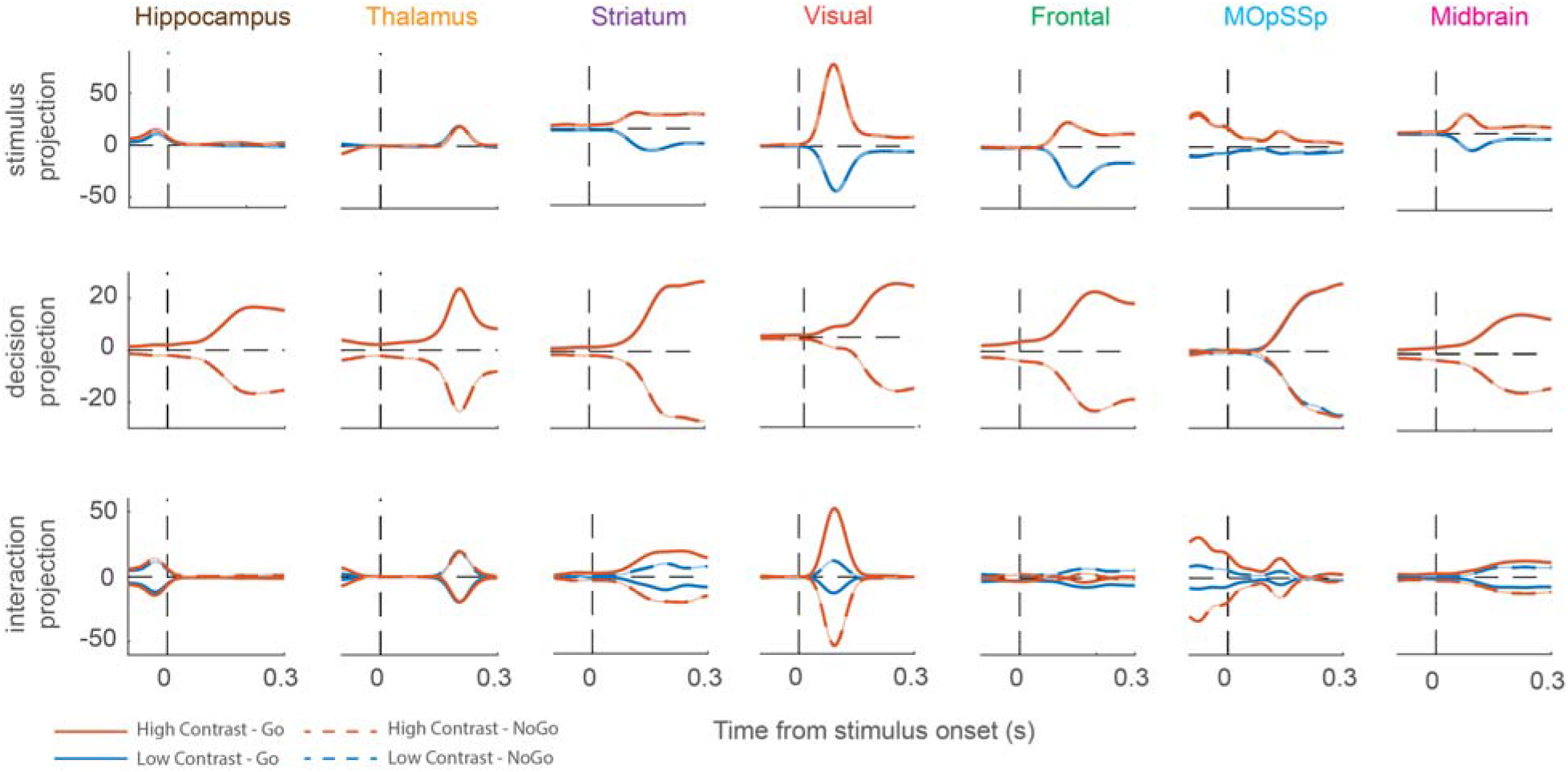
Demixed principal component analysis projections on task-related components. The figure shows the projection of population activity on stimulus (top row), decision (middle row), and stimulus-decision interaction (bottom row) components averaged over different components.

**Supplemental Figure 2.**
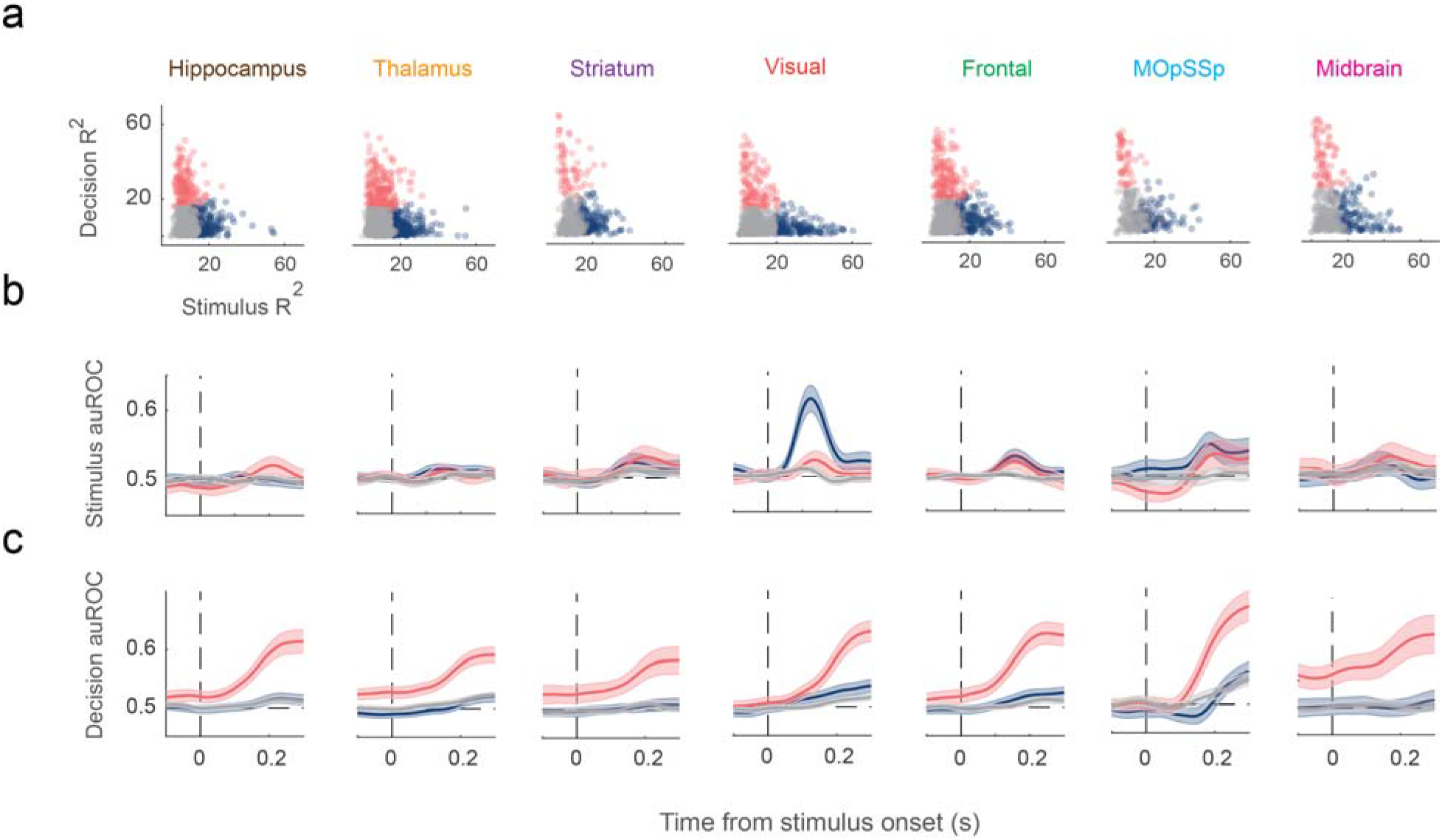
Separating neurons according to their response selectivity. a) Clustering of three neuron subtypes: stimulus (blue), decision (pink), and other (gray) based on demixed principal components. Panels b and c show the auROC validation of the three neuron subtypes. Stimulus neurons (blue) decode the stimulus much better than the other neuron subtypes. Decision neurons (pink) decode the decision (or detect probability) better than the other neuron subtypes. Shaded areas demonstrate CI.

**Supplemental Figure 3.**
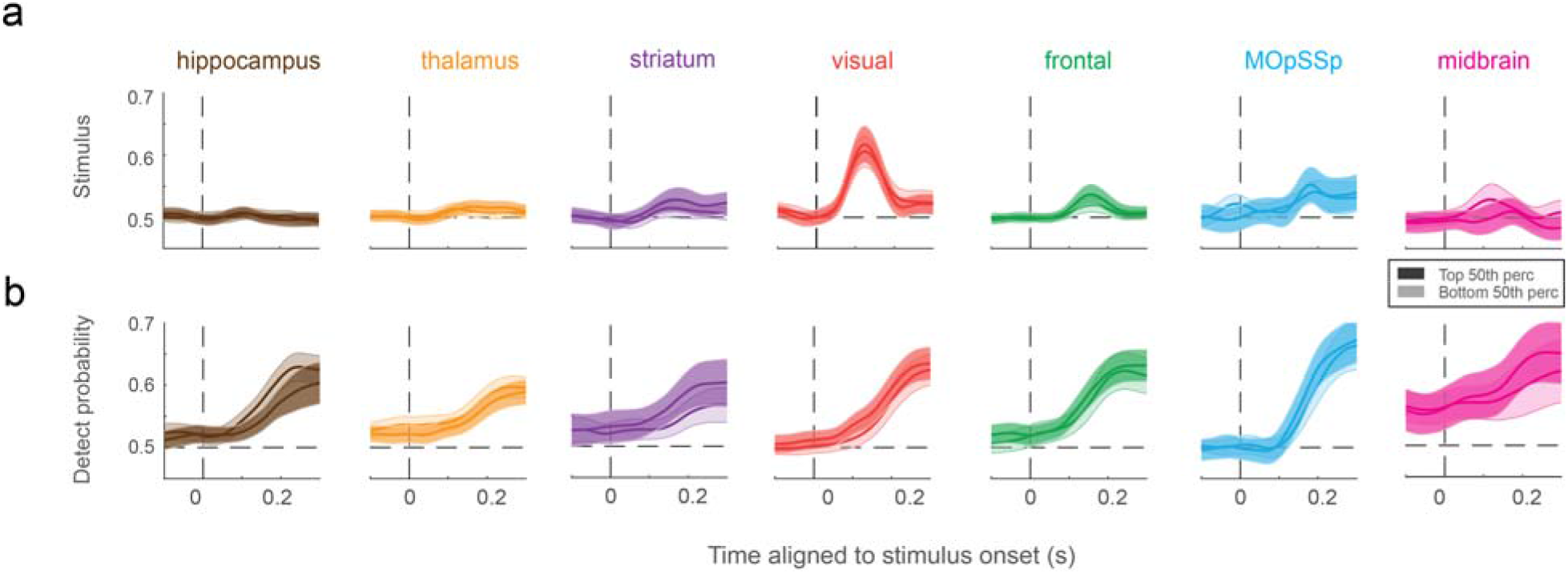
auROC timecourses for stimulus (panel a) and detect probability (panel b) for the short and long-timescale neurons. The curves represent the average of auROC values across neurons, and the shaded areas indicate the CI.

**Supplemental Figure 4.**
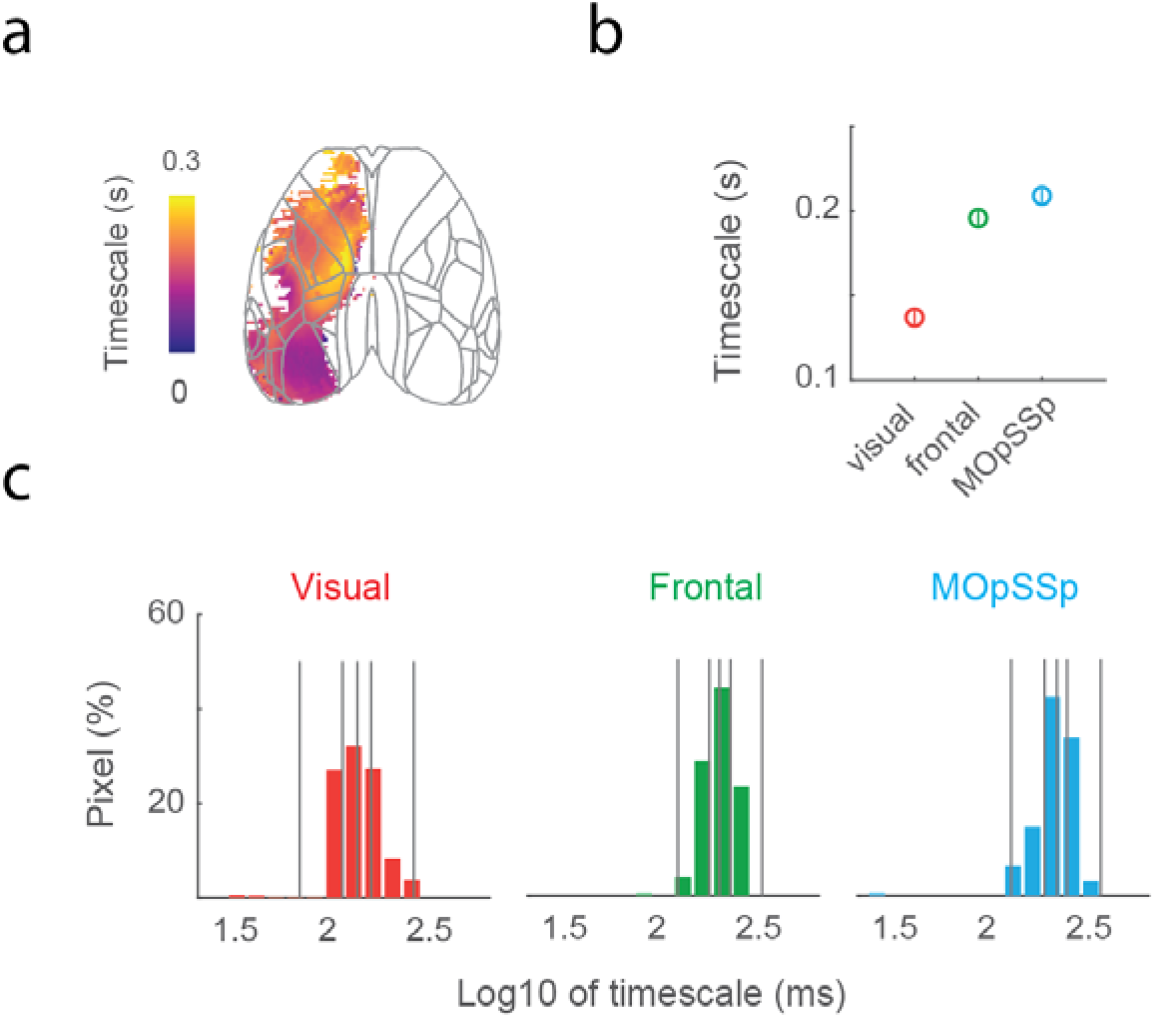
Timescale of the widefield calcium imaging data. a) Intrinsic timescale heatmap for the left hemisphere excluding pixels with poor timescale fits. b) The hierarchy of average timescales over pixels within each group area of the left hemisphere. c) Distribution of timescale within each brain region.

**Supplemental Figure 5.**
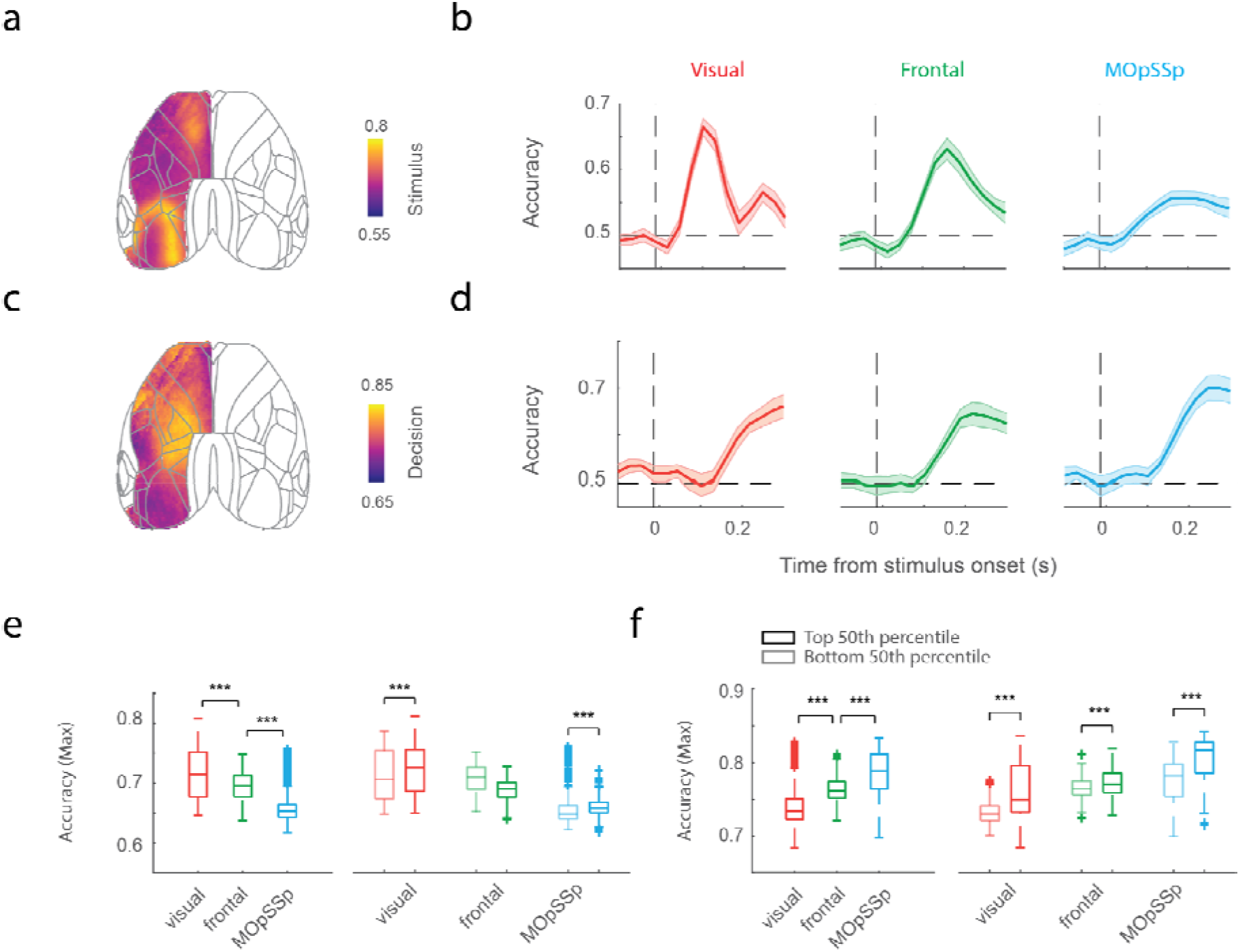
Stimulus decoding and detect probability on the fluorescence data. a,c) Heatmap of contra stimulus decoding accuracy and detect probability. b,d) Time course of the average decoding accuracy across pixels having valid timescale fitting for contra stimulus and detect probability. Shaded areas show CI. e) Max accuracy of contra stimulus for total valid pixels (left column) and median split of pixels based on the intrinsic timescale (right column). f) Max accuracy of detect probability for total valid pixels (left column) and median split of pixels based on the intrinsic timescale (right column). The sign ‘***’ symbolizes p-value < 0.001 in the Wilcoxon rank-sum test corrected for multiple comparisons.

## Data availability

All neural and behavioral data analyzed in this study are available at https://figshare.com/articles/steinmetz/9598406.

## Author contribution

M.M.G. conceptualization. E.I., A.H, and M.M.G. performed the analyses. E.I., A.H., and M.M.G. contributed to the methodology. E.I., A.H, and MMG carried out visualization. E.I., A.H., S.R. S.W.E., and M.M.G interpreted the results. E.I., A.H., S.R., and M.M.G. wrote the paper. S.R. and S.W.E review and editing, M.M.G. supervised the project.

## Competing interests

The authors declare no competing interests.

## Notes

### Competing Interest Statement

The authors have declared no competing interest.

